# Using cortical neuron markers to target cells in the dorsal cochlear nucleus

**DOI:** 10.1101/2020.09.19.304550

**Authors:** Thawann Malfatti, Barbara Ciralli, Markus M. Hilscher, Steven J. Edwards, Klas Kullander, Richardson N. Leao, Katarina E. Leao

**Affiliations:** Hearing and Neuronal activity Lab, Brain Institute, Federal University of Rio Grande do Norte, Natal, Brazil; Institute for Analysis and Scientific Computing, Vienna University of Technology, Vienna, Austria; Unit of Developmental Genetics, Department of Neuroscience, Uppsala University, Uppsala, Sweden

**Keywords:** Auditory system, channelrhodopsin, archaerhodopsin, unit, dorsal cochlear nucleus, ventral cochlear nucleus, CaMKIIα, Chrna2

## Abstract

The dorsal cochlear nucleus (DCN) is the first auditory region that integrates somatosensory and auditory inputs. The region is of particular interest for auditory research due to the large incidence of somatic tinnitus and increased aberrant activity in other forms of tinnitus. Yet, the lack of useful genetic markers for in vivo manipulations hinders the elucidation of the DCN contribution to tinnitus pathophysiology. In this work, we assessed whether adeno-associated viral vectors (AAV) containing the calcium/calmodulin-dependent protein kinase 2 alpha (CaMKII*α*) promoter and our mouse line of nicotinic acetylcholine receptor alpha 2 subunit (Chrna2)-Cre can be used to target specific DCN populations. The CaMKII*α* promoter is usually applied in studies of principal neurons of neo and paleocortex while Chrna2-cre mice express Cre recombinase in cortical dendrite inhibiting interneurons. We found that CaMKII*α* cannot be used to specifically target excitatory fusiform DCN neurons. EYFP expression driven by the CaMKII*α* promoter was stronger in the fusiform layer but labelled cells showed a diverse morphology indicating that they belong to different classes of DCN neurons. Light stimulation after driving Channelrhodopsin2 (ChR2) by the CaMKII*α* promoter generated spikes in some units but firing rate decreased when light stimulation coincide with sound presentation. Expression and activation of eArch3.0 (CaMKII*α* driven) in the DCN produced spike inhibition in some units but, most importantly, sound-driven spikes were delayed by concomitant light stimulation. We explored the existence of Cre+ cells in the DCN of Chrna2-Cre mice by hydrogel embedding technique (CLARITY). There were almost no Cre+ cell bodies in the DCN; however, we observed profuse projections arising from the ventral cochlear nucleus (VCN). Anterograde labeling Cre dependent AAV injected in the VCN revealed two main projections: one arising in the ipsilateral superior olive and the contralateral medial nucleus of the trapezoid body (bushy cells) and a second bundle terminating in the DCN, suggesting the latter to be excitatory Chrna2+ T-stellate cells). Stimulating ChR2 expressing terminals (light applied on the DCN) of VCN Chrna2+ cells increased firing of sound responding and nonresponding DCN units. This work shows that molecular tools intensively used in cortical studies may be useful for manipulating the DCN especially in tinnitus studies.

## 1 Introduction

The dorsal cochlear nucleus (DCN) of the auditory brainstem is the first integrator of auditory and multisensory signals and has been pointed as a key structure in tinnitus physiopathology (Kaltenbach et al., 2005; Tzounopoulos, 2008; Baizer et al., 2012). Cells in the DCN receives direct or indirect (e.g. relayed by the ventral cochlear nucleus - VCN) sound input onto different cell populations in a layer arrangement. The most cell-populated DCN field is the fusiform cell layer formed by excitatory fusiform cells intercalated with interneurons (Oertel and Young, 2004). An interesting aspect of the DCN is its architectural similarity to the cerebellum (Devor, 2000) that is thought to be responsible for integrative processing (e.g. sound/somatosensory, Oertel and Young, 2004).

Abnormal sensory integration in the DCN is clinically relevant due to the prevalence of temporo-mandibular tinnitus (Levine, 1999; Grossan and Peterson, 2017). Other forms of mechanical tinnitus are also attributed to aberrant activity in the DCN (Han et al., 2009). Also, a large number of studies have shown altered synaptic and intrinsic cellular properties within the DCN circuit relating to noise-induced tinnitus (reviewed by Shore et al., 2016) yet tinnitus treatments to date do not specifically target this region. The ventral cochlear nucleus (VCN) can also contribute to noise-induced tinnitus (Kraus et al., 2011; Coomber et al., 2015). Aberrant activity in any cochlear nucleus subregions can trigger upstream changes as cochlear nucleus neurons relay auditory signals to higher areas of the auditory pathway (Kraus et al., 2011; Coomber et al., 2015). Abnormal activity from auditory cortex and inferior colliculus can also produce downstream alterations in the DCN as its cells receive feedback through descending auditory fibers (Winer and Prieto, 2001; Milinkeviciute et al., 2016). Despite its physiological importance and its well accepted role in tinnitus, the contribution of specific DCN populations to hearing and tinnitus pathophysiology are largely unknown.

Due to its variety of cell types and its cerebellum like structure, DCN circuit studies could benefit from identifying key neuronal markers (Hilscher et al., 2017). The Calcium/calmodulin-dependent protein kinase 2 alpha (CaMKII*α*) promoter is widely used for targeting cortical pyramidal cells. Immunohistochemical data from rats has shown CaMKII*α* expression in the DCN molecular and fusiform cell layers (Ochiishi et al., 1998). In mice CaMKII*α* RNA is widely distributed in the fusiform layer (Lein et al., 2007). Hence, viral vectors to express of reporter or optogenetic proteins by CaMKII*α* promoter may be applied to DCN manipulation. Cortical interneuron markers could also be used to tag DCN cells. The calcium buffer protein parvalbumin (PV) is used for targeting fast spiking interneurons in studies of the hippocampus/neocortex (Kawaguchi and Kondo, 2002; Courtin et al., 2013) and PV expression specific to inhibitory neurons have also been described in some subcortical nuclei (Unal et al., 2015). In the DCN, PV is distributed across layers without population specificity. Moreover, somatostatin expression (found in dendrite targeting cortical/hippocampal interneurons) does not appear to follow a layer/cell specific expression (Lein et al., 2007), similar to the cortex/hippocampus where PV and somatostatin expression can be quite promiscuous (Kawaguchi and Kondo, 2002; Mikulovic et al., 2015). Recently, the nicotinic acetylcholine receptor alpha 2 subunit (chrna2) has been described as a marker for highly specific interneuron populations (CA1 oriens-lacunosum moleculare cells LM in the ventral hippocampus or L5 Martinotti cells in the neocortex; Leão et al., 2012; Hilscher et al., 2017). A Chrna2-cre mouse line was developed and evaluated, and significantly leaped the study of dendritic targeting interneuron populations (Leão et al., 2012; Enjin et al., 2017; Hilscher et al., 2017). Cre+ cells in Chrna2-cre mice seem to belong to single populations in several subcortical nuclei (Siwani et al., 2018). Expression of Chrna2 in the DCN is not described but a glance in whole brain imaging (clarity) evince almost no cell body expression in the DCN with Cre+ cell clusters in its vicinity and Chrna2+ positive axonal terminals that profusely target the DCN (Mikulovic et al., 2018; Siwani et al., 2018).

Here, we test if adeno-associated viral vectors (AAV) with the CaMKII*α* promoter can be used for manipulating DCN circuits in vivo. AAV encoded optogenetic protein expression and light stimulation paired with brief sound presentation was used to functionally identify cells and assess the effect of optical depolarization/hyperpolatization in input/output functions in CaMKII*α*+ neurons in combination with brief sound stimulation. Lastly, we examined how activation of Chrna2+ cells innervating the DCN modulate fusiform cell function.

## 2 Results

### 2.1 Auditory brainstem responses are normal during optogenetic excitation of CaMKII*α*-ChR2 positive DCN neurons

We first tested whether the Ca^2+^/Calmoduline kinase 2*α* (CaMKII*α*) promoter can be used to control subpopulations of DCN neurons. We injected viral vectors (rAAV5/CamK2-hChR(H134R)-eYFP or control rAAV5/CaMKIIa-eYFP) into the DCN for expression of ChR2 and/or enhanced yellow fluorescent protein (eYFP) under the control of the CaMKII*α* promoter of 1-2 month old C57Bl/6J mice. Four weeks later local protein expression in the DCN was examined, and showed strong eYFP signal in the vicinity of the injection site with a spread to both superficial and deeper layers of the DCN for both control and ChR2 vectors (Figure 1A, center and right, respectively). eYFP showed strong membrane expression of soma and neurites of DCN neurons for both ChR2 and control constructs, especially in cells with elongated somas perpendicular to the DCN edge with thick basal dendrites spreading towards the molecular layer (possible fusiform cells, Figure 1B, arrows). Smaller neuronal somas were also labeled with eYFP in the fusiform and deeper layers, as well as several large neuronal somas of the deep layer of the DCN with dendrites stretching along the internal edge of the DCN (possible giant cells, Figure 1C, arrow). This shows that the CaMKII*α* promoter is not specific for DCN excitatory neurons, although it seems to indeed label excitatory fusiform-like cells and giant cells with strong membrane expression.

**Figure 1:**
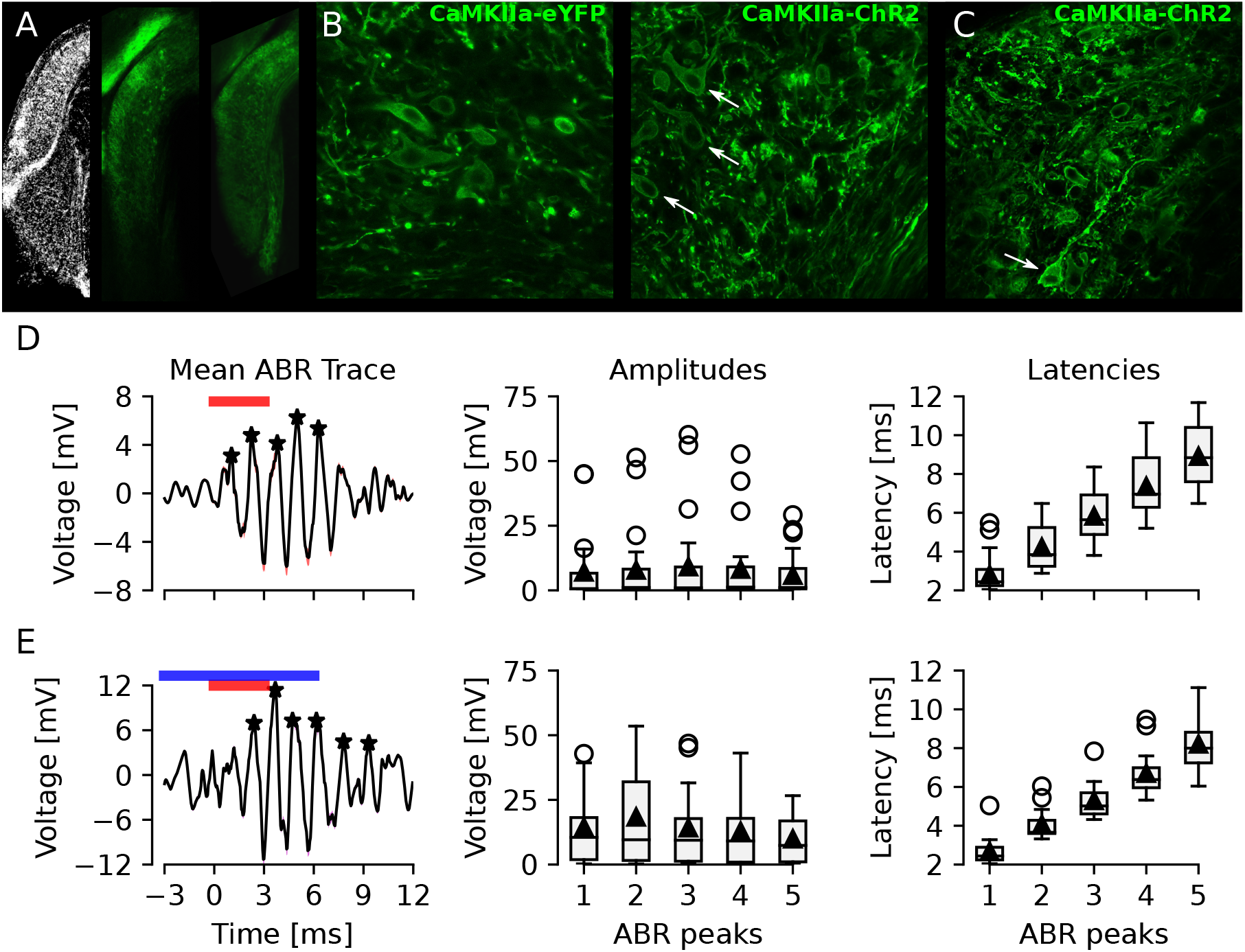
CaMKII*α*-ChR2-eYFP positive neurons in the DCN and normal auditory brainstem response in injected animals. **A)** Image of coronal brainstem sections with the DCN and VCN highlighted after DAPI nuclear staining (left), control CaMKII*α*-eYFP (center) and CaMKII*α*-ChR2-eYFP expression (right). **B)** High magnification confocal images showing several elongated horizontal somas (white arrows, possibly fusiform cells) labeled with membrane expression of eYFP. **C)** Another high magnification example of CaMKII*α*-ChR2-eYFP labeling of the DCN. Two possible giant cells are in the deep layer (white arrow). Lateral is left and ventral is down for all images. **D-E)** ABR waveforms recorded using electrodes lowered into the DCN in response to sound **(D)** and sound+light **(E)** stimulation protocols. Left, mean (black line) and SEM (red shadow) ABR traces (n=13), with detected peaks marked with black asterisks. Center, group amplitude of the first 5 ABR peaks. Right, group latency of the first 5 ABR peaks.

### 2.2 Optogenetic excitation of CaMKII*α*-ChR2 positive DCN neurons is decreased by concomitant sound stimulation

Next we wanted to examine how DCN units respond to optogenetic modulations using the CaMKII*α* promoter for expression of channelrhodopsin2 within the DCN. Mice previously injected with CaMKII*α*-ChR2 were anesthetized and fitted to a stereotaxic frame and a silicone 16-channel electrode was lowered vertically into the DCN (Figure 2A). A total of 224 isolated units were identified, of which 148 were excluded for not responding to neither sound nor light stimulus (Supplementary Table S2). From the remaining 76 units (n= 8 mice) 71% (54/76) responded to sound stimulation (3ms, 80dB, 5∼15kHz noise pulses presented at 10Hz) and response to sound was quantified and visualized by peristimulus time histograms (PSTHs; Figure 2B and E). Blue light stimulation (473nm, 10ms duration, at 10Hz with intensity of 5mW/mm^2^ at fiber tip) delivered by a glass optic fiber to the DCN (Ø200*μ*m, inserted in a 45° angle from the contralateral side; Figure 2A), elicited increased firing of units immediately following blue light stimulation (Figure 2C and E). We found 25% (19/76) units responding to light stimulation.

**Figure 2:**
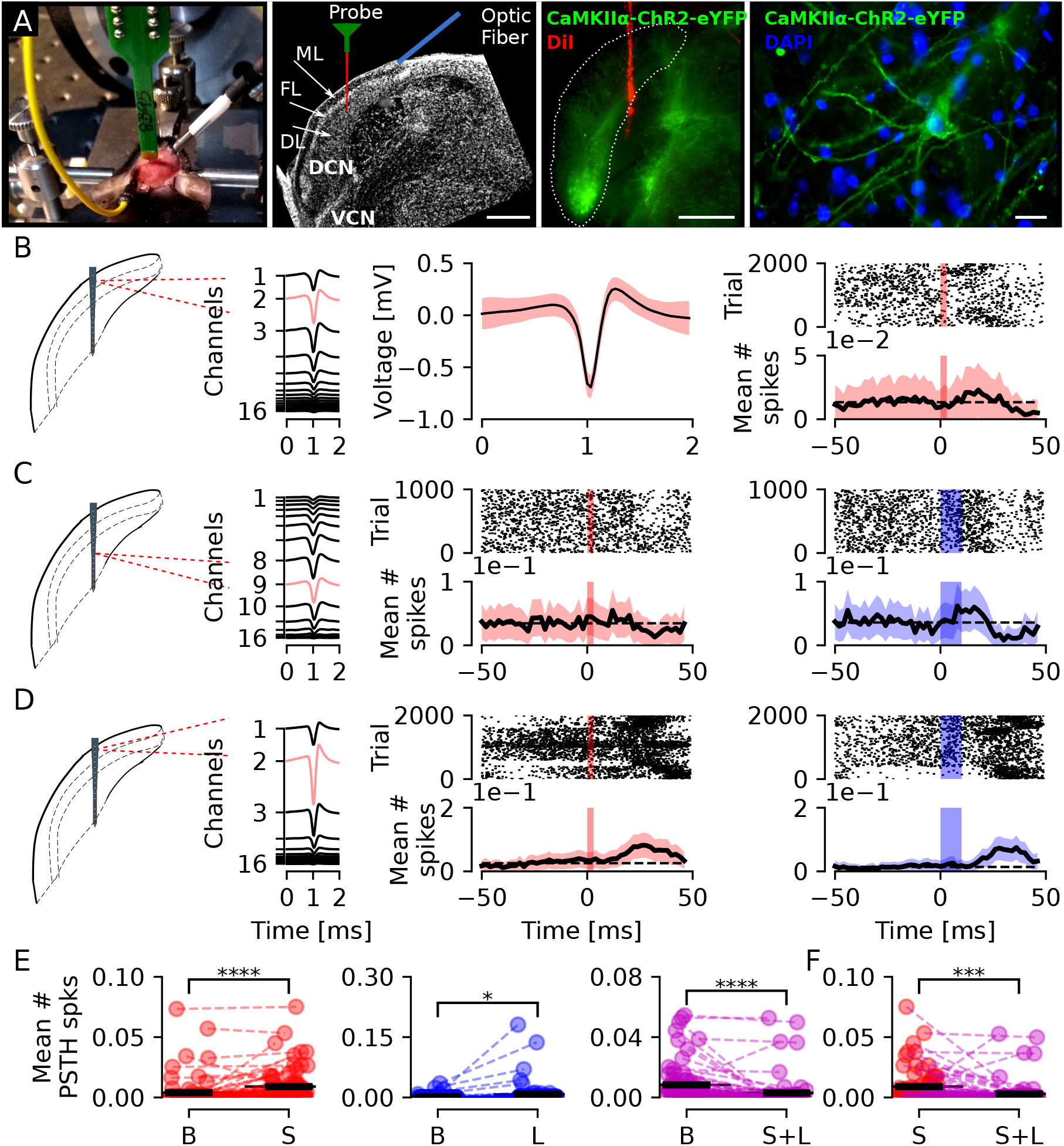
Activation of CaMKII*α*-ChR2-eYFP expressing neurons in the DCN during sound stimulation can decrease unit firing. **A)** Left, photography of a mouse in the recording setup and schematic representation of recording electrode and optic fiber for blue light stimulation in a DAPI labeled section showing the DCN sub-regions (ML-molecular layer, FL-fusiform cell layer, DL-deep layer). Center, confocal image showing an example of eYFP expression in the dorsal region of the DCN with the probe tract colored by DiI. Scale bar: 100*μ*m. Right, example of a DCN neuron expressing eYFP along the somatic membrane and proximal dendrites. Scale bar: 20*μ*m. **B)** Example of a unit with its waveform shown at higher magnification (center), that responded to sound stimulation by briefly (∼20ms) increasing its firing. **C)** An example of a unit that does not respond to sound stimulation but increases firing in response to blue light stimulation. **D)** Example of a unit responding to sound (center) and blue light (right) stimulation. **E)** Group mean number of spikes for all units (n=76) showing a significant increase after sound (S) comparing to baseline (B; left) or blue light (L; center) stimulation (p = 2.6e-5 and 0.016) and a significant decrease after concomitant sound and light stimulation (S+L; right; p=4.2e-5). **F)** Group mean number of spikes for all units (n=76) after sound stimulation is significantly higher than after concomitant sound and light stimulation (p=3.8e-4).

Out of these, 58% (11/19) responded exclusively to light stimulation, 26% (5/19) responded to either sound or light stimulation (Figure 2D), and 16% (3/19) responded only to a combination of sound and light stimuli.

The majority of units, 86% (65/76), had firing rates from 0.05∼6.46Hz (average 1.5 ± 0.19) while a smaller proportion of units recorded had higher firing rates (14%; 11/76), from 8.83∼72.43Hz (average 26.76 ± 5.81 Hz; Supplementary Table S2). Including all 76 responding units, averaged over 1.5min including stimulus epochs, and comparing to specific stimuli showed a significant increase in response to sound (p=2.6e-5; Figure 2E left) or light (p=0.016; Figure 2E center) but, interestingly, a decrease in response to concomitant sound and light stimulation (p=4.2e-5; Figure 2E right). Also, the mean number of spikes in response to sound were significantly higher than to concomitant sound and light (p=3.8e-4; Figure 2F). This shows that optogenetic excitation using ChR2 can increase firing of DCN units, even in units not directly responding to sound or light, but controversially, presence of simple sounds during optogenetic stimulation can decrease the over all unit firing.

### 2.3 Inhibiting CaMKII*α*-eArch3.0 positive neurons in the DCN delay response to sound and bidirectionally affect units not responding to sound

We then tested if we could inhibit CaMKII*α*-eArch3.0 positive DCN neuron response to sound using the outward proton pump eArch3.0. We injected adult (1 month) wild-type C57Bl/6J mice unilaterally with Archaerhodopsin-containing viral vectors (CaMKII*α*-eArch3.0-eYFP) and 4 weeks later extracellular activity in the DCN was recorded. Units were recorded in response to short sound pulses (3ms, 80dB, 5∼15kHz noise pulses presented at 10Hz) and next using an optic fiber coupled to a green laser source (543nm excitation) we examined if response to sound could be abolished by concurrent green light stimulation (543 nm, 20s, repeated 5x with 10s interval, so it is concomitant to sound pulses). Out of 86 units isolated (n=4 mice), we found that 17/86 (20%) units responded to stimulation (Figure 3D, left), from which 12/17 (71%) responded to sound stimulation (Figure 3E, left). The most striking finding was that instead of inhibition of responses to sound we found several types of unit responses being delayed during green light stimulation (Figure 3A-B). 7/17 (41%) units that sharply responded to sound showed a delayed response to sound with mean latency of 19.1 ± 1.22ms when the DCN CaMKII*α*-eArch3.0 cells were inhibited during sound stimulation (Figure 3A-C and E, right). The delayed response to sound continued strongly time-locked. This could suggest that some DCN CaMKII*α*-eArch3.0 positive neurons may be inhibitory, as green light stimulation could involve disinhibition of cartwheel cells, and complex spiking units are known to generate delayed responses in PSTH in the range of 10-40ms delay (Parham et al., 2000). Examining responses from the units isolated, the majority (10/12) of units responded to sound stimulation by increasing firing rate (Figure 3A-B), while 2/12 units decreased the firing rate in response to sound stimulation (Figure 3C) and both such responses were delayed in the presence of green light stimulation. Furthermore, 5/17 units responded exclusively to sound, while another 5/17 units responded exclusively to the combination of sound and green light (Figure 3D, right).

**Figure 3:**
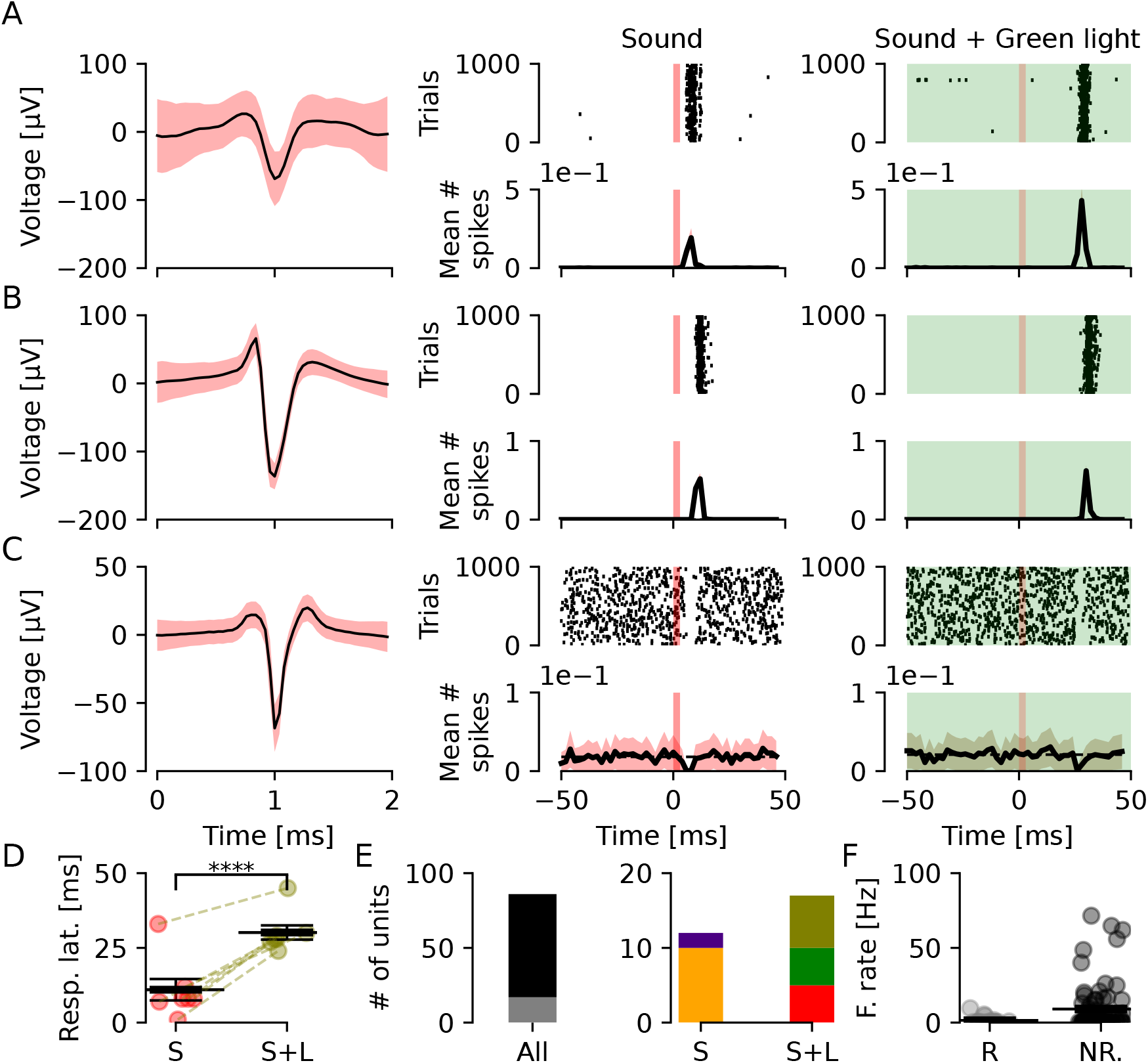
Inhibition of CaMKII*α*-eArch3.0 expressing neurons can delay unit sound responses. **A)** Example of a unit that responds to sound, while during inhibition of CaMKII*α*-eArch3.0 positive DCN cells this unit shows delayed excitation. **B)** Another unit showing distinct time-locked excitation in response to sound stimulation. This excitation is delayed by ∼20ms when CaMKII*α*-eArch3.0 positive DCN cells are inhibited. **C)** Example of a DCN unit with a negative response following sound stimulation. This pause in firing is delayed by CaMKII*α*-eArch3.0 positive DCN cells inhibition. **D)** Group latency in response to sound is significantly increased for all units responding to both stimulation (n=7, p=6.7e-6). **E)** Quantification of numbers of units. Left, out of 86 units, 17 (gray; 20%) responded to provided stimulation. Right, out of 12 units responding to sound, 10 (orange; 83%) increased and 2 (purple; 17%) decreased firing in response to sound. Out of 17 responding units, 5 (red, 29%) responded only to sound, 5 (green, 29%) responded only to sound and light combined, and 7 (olive, 41%) responded to both stimulation. **F)** Group firing rate of responding and non-responding units.

Examining firing rates of sound responding units and comparing to non-responding units showed that we targeted slow firing units responding to sound (1.5 ± 0.6Hz and 9.1 ± 2Hz for responding and non-responding units, respectively). Still, it is also important to examine the effect of DCN inhibition on units not responding directly to sound. We found, for the remaining 69/86 units that did not respond to stimulation, 39% (27/69) of units to decrease spontaneous firing rate under green light stimulation (Figure 4A and C). Out of the 27 units, 12 were high frequency firing (33.51 ± 6.3Hz) that decreased firing to 70% of the initial frequency (23 ± 6.68Hz) under green light stimulation, while 15 units were low frequency firing (2.19 ± 0.57Hz), decreased firing frequency by 58% (1.26 ± 0.48Hz) upon green light stimulation. On the contrary, 42 units increased firing frequency upon green light stimulation (Figure 4B and C) where 11 high frequency firing units increase in firing frequency to ∼ double the initial frequency (16.4 ± 4.38Hz to 34.11 ± 6.76 Hz) while 31 low firing units on average increased firing from 0.33 ± 0.11 Hz to 1.54 ± 0.29 Hz (Figure 4B and C). Overall, non-responding units that had low firing rate in response to sound showed a significant increase in firing rate (n=46/69 units, p=0.03). Also, CaMKII*α*-eArch3.0 inhibition caused a bidirectional effect, with 27/69 decreasing (p=4e-4) and 42/69 increasing (p=0.01) firing rate (Figure 4C). This highlights the complexity of the DCN and how precaution must be taken when attempting to decrease neuronal activity *in vivo* of the auditory brainstem using tools such as CaMKII*α*-eArch3.0.

**Figure 4:**
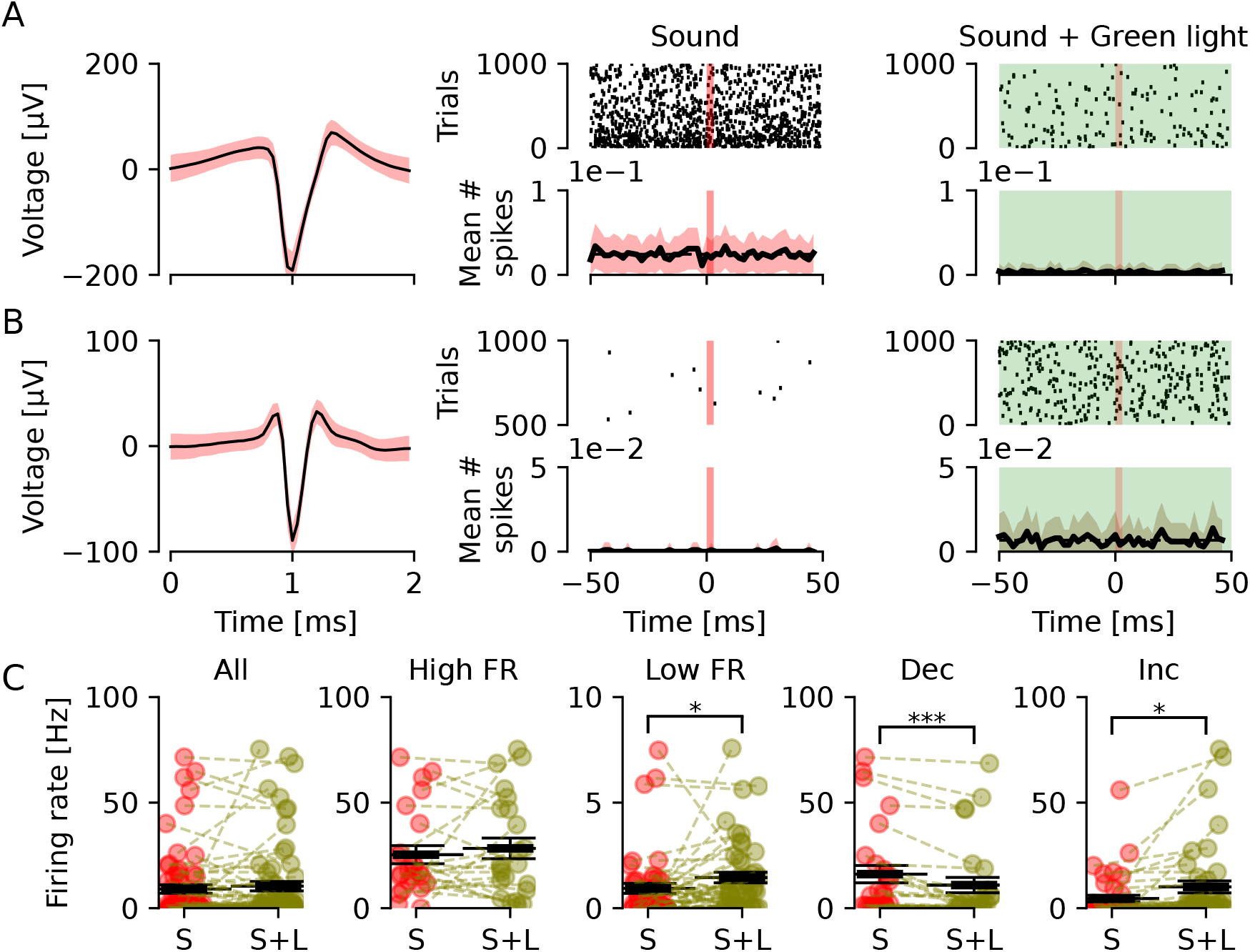
Inhibition of CaMKII*α*-eArch3.0 positive cells in the DCN can both increase and decrease excitation of DCN units. **A-B)** Extracellular unit recordings from the DCN of an anesthetized mouse previously injected with CaMKII*α*-eArch3.0-eYFP viral vectors under sound stimulation and concomitant sound and green light exposure, from two distinct units not responding to sound stimulation. Units are in the same DCN region (strongest signal at ∼-3.5mm DV), that have different waveforms and different baseline firing rates. Even though none of the units responded to sound stimulation, one of the units decreased firing rate **(A)** while and the other **(B)** increased its firing rate upon inhibition of CaMKII*α*-eArch3.0 DCN cells. **C)** Group firing rate of all non-responding units (n=69) under sound (red) or concomitant sound and green light (olive). Units were divided into high and low firing rate, and a significant increase in firing was found for low firing units (p=0.03). Units were also divided into units that decrease or increase firing rate comparing both stimulations, and a significant decrease and increase was found (p = 4.7e-4 and 0.01, respectively).

### 2.4 The Chrna2-cre transgenic line targets putative T-stellate cells and bushy cells of the VCN

To confirm that viral constructs do not cause any abnormalities in hearing we routinely extracted auditory brainstem response (ABR) waveforms from extracellular recordings. A high impedance electrode (16 channel single shank silicon probe, Neuronexus), placed in the DCN for recording unit responses in anesthetized mice, was used to isolate ABRs in response to sound or concomitant sound and optogenetic stimulation. The anatomical correlation between auditory brainstem structures and ABR peaks, where I corresponds to the auditory nerve; II, cochlear nuclei; III, superior olivary complex; IV and V, inferior colliculus (Henry, 1979) is useful for verifying an intact auditory brainstem system. We found all animals to display normal mean ABR waveforms (n=13) in response to 80 decibel sound pressure level (dBSPL) stimulation (Figure 1D). Also, ABR mean amplitude and latency was not affected by concomitant sound and light stimulation (Figure 1E). Together this show that the animal’s hearing at the stimulus intensity was not impaired by the viral vector injection procedure or by concomitant blue light stimulation.

A recent and interesting transgenic cre-line is the chrna2-cre mouse that targets specific interneuron populations in different brain and spinal cord regions (Leaõ et al., 2012; Mikulovic et al., 2015; Hilscher et al., 2017; Perry et al., 2015; Siwani et al., 2018). So far there are no reports of Chrna2-cre expression in auditory areas, so here we start by exploring Chrna2-cre expression in the cochlear nucleus and superior olivary complex. Heterozygous Chrna2-cre transgenic mice were crossed with homozygous tdTomato-lox reporter mice to visualize Chrna2 positive (Chrna2+) cells in the auditory brainstem (Figure 5 and 6A, Supplementary Video S1). We found nicotinic acetylcholine receptor *α*2 subunit (Chrna2) positive cells in the VCN, with dense projections of axons branching into the DCN as well as to the superior olivary complex (Figure 5 and 6A). Brains processed for CLARITY examination (Hilscher et al., 2017) also show Chrna2+ VCN cells (white) with some projections to the DCN (Supplementary Video S1). In order to identify the boundaries of DCN and VCN and examine any soma labeling in the DCN, slices of Chrna2-tdTomato animals were co-stained with DAPI (Figure 5C). Very few red labeled somas were identified in the DCN of Chrna2-tdTomato animals or in CLARITY images and did not relate to any particular region (Figure 5C). Next, Chrna2-cre mice were injected with cre-dependent ChR2 (Chrna2-cre/DIO-ChR2-eYFP) viral vector into the VCN, and the expression pattern was similar to the Chrna2-tdTomato (Figure 5D). Based on projection patterns we assume Chrna2+ cells of the VCN to comprise of both stellate and bushy cell subtypes (Figure 5 and 6A). Previously, T-stellate cells have been shown to respond to cholinergic agonists (have acetylcholine receptors), while D-stellate cells are insensitive to carbachol (Fujino and Oertel, 2001), therefore Chrna2-cre positive neurons projecting to the DCN (Figure 5E) are most likely T-stellate cells (Oertel et al., 2011). T-stellate cells also projects to the ipsilateral LSO (Oertel et al., 2011), which supports the strong labeling of ChR2 in the ipsilateral LSO (Figure 5D). The ipsilateral LSO is also labeled by VCN bushy cells as projections to the contralateral medial nucleus of the trapezoid body (MNTB, Figure 5F) as well as the ipsilateral LSO were apparent, thereby indicating that Chrna2-cre labels both globular and spherical bushy cells.

**Figure 5:**
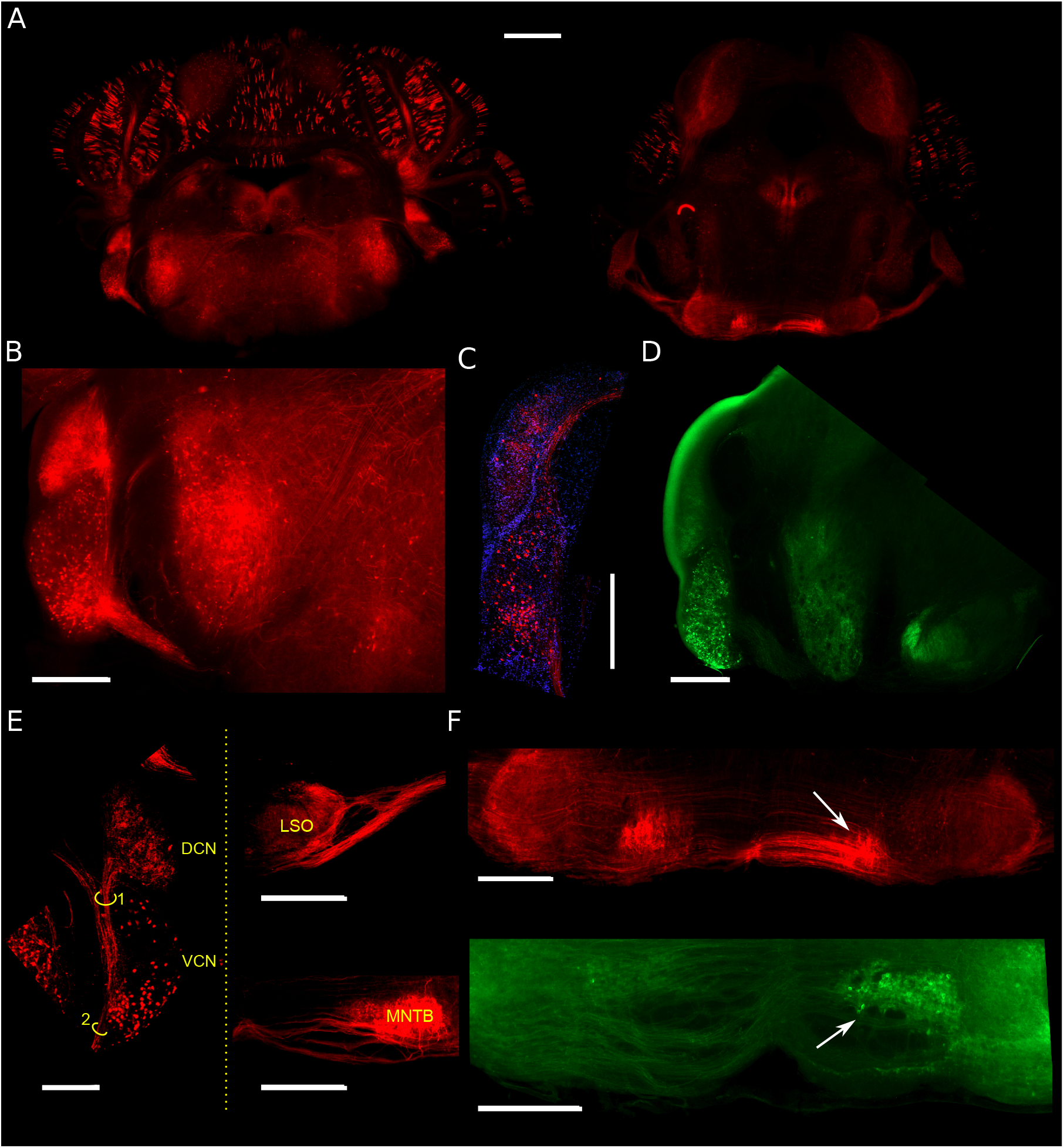
Confocal images showing tdTomato expression and Chrna2-cre/DIO-ChR2-eYFP expression in transversal brainstem sections from Chrna2^cre^/tomato^lox^ mice. **A)** Mosaic of images showing a coronal overview of Chrna2-tdTomato expression at cochlear nucleus (A) and medial nucleus of the trapezoid body (MNTB; B) anteroposterior coordinate. **B)** Zoom-in image showing tdTomato expression in the VCN and DCN. Red cell bodies can be clearly seen in the VCN area while red expression is more diffuse in the DCN, suggesting this area is showing dense axonal terminations from VCN T-stellate cells. **C)** Another example showing Chrna2-tdTomato expression overlayed with DAPI staining. **D)** Image showing unilateral expression of eYFP following local injections with cre-dependent ChR2 (Chrna2-cre/DIO-ChR2-eYFP) constructs in the VCN. The VCN contains strongly labeled cell bodies and the DCN shows diffuse green labeling. The strong green edge of the DCN is an artifact of the mounting medium. The ipsilateral S-shaped LSO is also strongly labeled by eYFP. **E)** Image of a coronal slice of a Chrna2^cre^/tomato^lox^ mice showing the cell bodies in the VCN and projections going up to the DCN. Highlighted bundles project to LSO (1) and MNTB(2). **F)** Top, zoom-in showing strong labeling of the MNTB and the lateral superior olive (LSO; arrow) suggests that also anteroventral VCN bushy cells are expressing the Chrna2 promoter. Bottom, a subsequent brainstem section (from the same injected animal from **D**) showing strong labeling of the contralateral MNTB (arrow). Scale bars: 1mm (**A**) and 500*μ*m (**B** and **C**).

**Figure 6:**
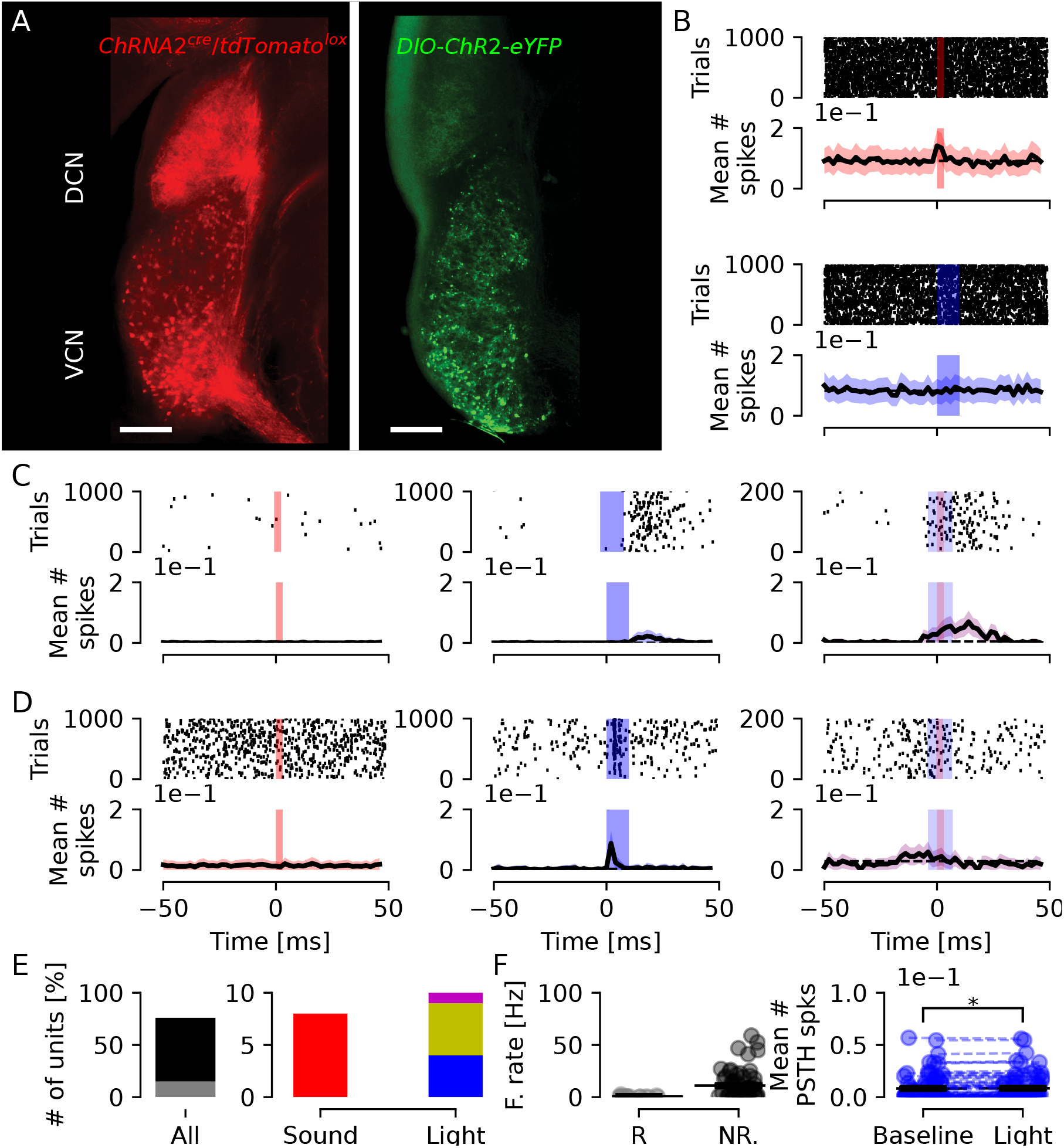
Chrna2-cre positive neurons of the VCN can be targeted to drive activity of DNC neurons. **A)** Left, confocal image showing tdTomato (red) expression in VCN cell bodies and strong axonal arborizations in the DCN fusiform and deep layers. Right, expression of ChR2-eYFP (green) in Chrna2-cre positive neurons of the VCN, with diffuse green innervation of the DCN. **B)** Example of a unit from the DCN that respond to sound but not light stimulation. **C)** Example of a unit from channel 10 that does not respond to sound stimulation, but responds to light stimulation (middle). In 200 trials of combined light and sound stimulation the unit responds to both stimuli, with what appears as an anticipation of sound. **D)** Example of a unit not responding to sound pulses but responds with high fidelity to light stimulation of the VCN. When sound and light stimuli were combined, the unit failed to respond. **E)** Quantification of number of units according to response. Left, out of 76 units, 15 (gray) responded to stimulation. Right, out of 15 responding units, 8 (red; 53%) responded to sound; 4 (blue; 27%) responded only to light; 5 (yellow, 33%) responded to both sound and light and only 1 (magenta, 7%) responded to light or concomitant sound and light stimulation. **F)** Left, group firing rate of responding and non-responding units. Right, group mean number of spikes showing a significant increase after light stimulation (n=76 units, p=0.026).

### 2.5 VCN Chrna2 positive neurons can generate indirect optogenetic activation of DCN neurons

To investigate if optogenetic control of putative T-stellate cells can excite DCN neurons, Chrna2-cre mice were unilaterally injected with cre-dependent ChR2 (Chrna2-cre/DIO-ChR2-eYFP) into the VCN generating labeling of diffuse fibers spreading in the deep layers of the DCN (Figure 6A). To investigate if DCN cells can be indirectly excited by light stimulation of ChR2-eYFP expressing Chrna2+ cells of the VCN, we recorded unit responses in the DCN from anesthetized mice while stimulating with blue light (10ms, 473nm light pulses at 10 Hz) with an optic fiber placed in a 45° angle into the VCN. We first analyzed if units responded to sound stimulation. Out of 76 units extracted, only 15 units (n=8, 6 and 1, from three mice respectively) responded to sound and/or light stimulation (Figure 6B and E). For units not responding to sound, we found examples of a DCN unit with response to VCN light stimulation that was prolonged and appeared increased in the presence of sound (Figure 6C). Also, one DCN unit with highly temporally precise responses to VCN light showed a loss of response in the presence of sound (Figure 6D). In summary, 4/15 units responded only to light stimulation, 2/15 units responded only to sound stimulation (Figure 6B and E), and 13/15 units responded to light or combined light and sound stimulation (Figure 6C and E). Excitation of VCN Chrna2+ cells significantly increased firing rate in DCN units, both responding and not responding to sound (Figure 6F). Still, the identity of these different units has to be further investigated but highlights that the Chrna2-cre line has potentials for auditory research.

## 3. Discussion

There is a large variety of genetic tools for dissecting the role of specific neuronal populations. The vast majority of these tools are transgenic mice expressing a reporter protein or a recombinase. Attempts to engineer viral vectors with population specific promoters have not produced useful tools for cell specificity. Currently widely used viral vector/promoter based gene expression in neuroscience are limited to hSyn (synapsin) for all neurons, CamK2a for cortical pyramidal cells, mDLx for interneurons, GFAP for glia and a cohort of generic (any cell) promoters. In regard to transgenic mice (e.g. expressing cre recombinase under the control of specific gene expression), there is a large library of strains for targeting specific interneuron populations in the cortex or single neurotransmitter systems in subcortical nuclei like SERT-cre (serotoninergic), DAT-cre (dopaminergic), Chat-cre (cholinergic) etc. However, there is, to our knowledge, no Cre line or promoter based vector for specific tagging of auditory neurons, especially those residing in the brainstem.

In this work, we first show that the CaMKII*α* promoter targets DCN neurons of different morphology, not only putative fusiform cells. Next, we show that excitation of CaMKII*α*-ChR2 positive DCN cells during sound stimulation generates normal ABRs, and does not disrupt hearing pathways. Still, we show that optogenetic excitation of CaMKII*α*-ChR2 positive DCN cells modulates DCN unit firing rate, and that such light stimulation is sensitive to concurrent sound stimulation. When aiming to inhibit DCN activity we found that inhibiting CaMKII*α*-eArch3.0 positive DCN cells can delay response to sound instead of decreasing DCN units firing rate.

More so, in units not responding directly to sound, CaMKII*α*-eArch3.0 inhibition of the circuit could bidirectionally alter firing rate of units. Lastly, we show that the Chrna2-cre line strongly labels cell bodies in the VCN and that these cells are putative T-stellate cells, based on literature and projection patters; and bushy cells, based on innervation to the MNTB and LSO. Furthermore, Chrna2-cre/DIO-ChR2 excitation in the VCN could increase firing of a small number of DCN units and this excitation appeared temporally precise. Still, this connectivity needs to be further characterized both pre and post synaptically.

Our experiments were performed in C57BL/6J mice as these animals are commonly used for genetic manipulations with optogenetic tools. It is known that the C57 mouse strain has a point mutation in the *cdh23* gene (Noben-Trauth et al., 2003) and thereby suffers a progressive high-frequency hearing loss after approximately 3 months of age. Therefore we perform all experiments in mice around 2 months of age, with viral constructs injected at 1 month of age and allowed 3-4 weeks for adequate protein expression before optogenetic experiments. Still, it is important to confirm that animals indeed can hear the tested frequencies used in experiments. Here, extracted auditory brainstem responses showed typical ABR peaks indicating neuronal responses to sound at several levels of the auditory brainstem, and thereby intact hearing to the tested stimuli. Since extracellular recordings can pick up ABR signals, ABR protocols can be useful to routinely add to single or multi unit recordings in the DCN especially when using older mice on a c57BL/6J background. Recently, new cre-lines expressing channelrhodopsin on a CBA background, with preserved hearing throughout adult life, has been developed (Lyngholm and Sakata, 2019). Lyngholm et al. (2019) show that CBA mice expressing ChR2 coupled to the parvalbumin promoter could excite cortical narrow spiking neurons (inhibitory interneurons) upon light stimulation, and inhibit broad spiking units as effectively as in mice with a C57 background (Lyngholm and Sakata, 2019). On the other hand, a benefit of using the c57BL/6J mouse line is that it is more susceptible to acoustic overexposure than other strains (Willott and Erway, 1998; Davis et al., 2001) and thereby a suitable animal model for studying noise-induced tinnitus.

While several studies have applied general promoters to achieve optogenetic control of the DCN (Shimano et al., 2013; Darrow et al., 2015; Hight et al., 2015) there is still a lack of studies showing subpopulation control of the DCN *in vivo*. The CaMKII*α* promoter is often used for targeting excitatory neurons in the neocortex and hippocampus (Wang et al., 2013) but here we found that the CaMKII*α* promoter, for the viral constructs tested, would target morphologically different cell types within the DCN. This is in agreement with studies of the olfactory bulb, where CaMKII*α*-GFP positive neurons co-localize with GABA immunoreactivity (Wang et al., 2013). Still, to decrease activity of DCN excitatory neurons would be highly interesting for alleviating tinnitus. In slice preparations, vesicular glutamate transporter 2 (VGluT2) transgenic mice has already been used for targeting and controlling DCN fusiform cell firing (Apostolides and Trussell, 2013a). Inhibitory DCN activity have been investigated using glycine transporter 2 (GlyT2-cre mice) for controlling DCN cartwheel cell firing *in vitro* (Apostolides and Trussell, 2013b; Lu and Trussell, 2016). Also the GABA/glycine transporter (VGAT) promoter has been used to excite inhibitory interneurons to study inhibitory neurotransmission in slices of the VCN (Xie and Manis, 2014) for example.

Attempting to silence DCN units responding to sound, using light stimulation of CaMKII*α*-eArch3.0 expressing neurons showed that units could be inhibited using eArch3.0, but also that green light exposure generated a distinct delay in response-onset to sound. The delay was consistently around 20ms suggesting polysynaptic activity, possibly from the recruitment of complex spiking cartwheel cells that can respond with 20ms delay to pure tone stimulation (Parham et al., 2000). Also PSTH of unidentified neurons with a 40ms delay between initial and basal firing have been reported for guinea pigs (Robertson and Mulders, 2018). Our experiment could not identify the type of unit responsible for this delay, but it shows that silencing CaMKII*α*-eArch3.0 expressing neurons is not enough to disrupt sound generating activity in the DCN circuit. Some studies have pointed to technical problems when using the proton pump eArch3.0, as it may affect intracellular pH of presynaptic membranes and promote neurotransmitter release if light is applied continuously for several minutes (Mahn et al., 2016). Mahn et al. (2016) showed that 5 min of continuous eArch3.0 activity significantly increased the EPSC rate (Mahn et al., 2016). Thereby, eArch3.0 may not be the most appropriate tool for inhibiting neurons of the DCN for longer time periods. Here we found that applying green light stimulation in blocks of 20s was sufficient to alter temporal coding of some units and silence others. An advantage of green light stimulation, compared to blue, is that green wavelength light can penetrate deeper into tissue without being scattered. For example, it has been shown in a modeling study that green light penetrates skin tissue twice as deep as blue light (Ash et al., 2017). Thereby, the green light applied here should be adequate for illuminating the DCN, and not be the reason for failed neuronal silencing in units showing delayed sound response. Still, the placement of the recording electrode could influence our findings as we are only sampling local neurons according to the probe location. As we describe in methods, we adjusted our coordinates to the animal’s skull size and recorded at three different depth to cover an as large as possible region of the DCN for each animal, without inserting the probe at multiple ML/AP locations or aspirating the cerebellum, as done in other rodent studies (Kaltenbach and Zhang, 2007; Shore et al., 2007; Finlayson and Kaltenbach, 2009; Koehler et al., 2010; Dehmel et al., 2012; Manzoor et al., 2012). Furthermore, spontaneously firing neurons that decreased or increased firing upon green light stimulation were recorded all along the probe, showing that they were not from any specific DCN layer. An interesting finding from CaMKII*α*-eArch3.0 experiments was that many neurons not responding directly to brief sound of 5-15kHz were indirectly affected by silencing CaMKII*α*-eArch3.0 positive neurons in the DCN. Also, that inhibiting neurons expressing CaMKII*α*-eArch3.0 can be used to modulate both high and low frequency firing neurons not responding directly to sound, and that this modulation was bidirectional. This suggests that CaMKII*α* expressing neurons can modulate excitation/inhibition ratios to some extent. Here, stimulating the DCN with longer sound at additional frequencies would clarify what type of units respond by decreasing or increasing spike activity when inhibiting CaMKII*α* positive DCN neurons.

A limitation of our study is that we did not assess the best frequency of units. Thereby the sound stimulus will not display full firing potential nor specific firing patterns (such as pauser, onset, build-up units). However, as light stimulation also was brief, it allows for more direct comparison between modulation of unit activity in responses to sound or after exciting DCN units with light. We speculate that the decrease in firing when stimulating CaMKII*α*-ChR2 positive neurons with blue light pulses during sound stimulation could be part of motifs of feed-forward inhibition (Roberts and Trussell, 2010); or light masking (Hernandez et al., 2014), where the cell do not respond to sound because it is in refractory period after responding to light stimulation.

Our work also show for the first time that the Chrna2-cre line targets two different cell population of the VCN; both putative T-stellate cells and bushy cells of the VCN. Using cre-dependent viral constructs we could excite DCN units that did not respond directly to sound, but responded temporally precise to light stimulation of the VCN, suggesting that specific circuits may be targeted using these animals. We found no disruption of hearing upon optogenetic stimulation, but specifically altered network activity compared to brief sound stimulation, including delayed responses and disinhibition of activity. Interestingly, we also found altered spiking activity for units not responding directly to sound. Furthermore, stimulation of bushy cells may especially be useful for studies of the calyx of Held presynaptic release and/or sound localization studies using the Chrna2-cre line. Together, these results opens up for more detailed control of DCN circuit output in vivo and novel tools for studying tinnitus mechanisms.

## 4. Conclusion

Optogenetic stimulation of CaMKII*α* positive DCN neurons or Chrna2 expressing VCN neurons can be used to manipulate unit firing of the DCN and to modulate response to sound. In general, we found the CaMKII*α* and Chrna2 promoter to be interesting tools as smaller and more specific network activity modulation was achieved compared to sound stimulation.

## 5. Methods

### 5.1 Mice

Male C57Bl/6J mice, Chrna2-cre or Chrna2-cre mice crossed with the reporter line Ai14 tdTomato (Chrna2-Cre/R26^tom^) mice age P21-P75 (n=14) were used in this study. All animal procedures were approved by the Federal University of Rio Grande do Norte Ethical Committee in Use of Animals (CEUA - protocol number 051/2015) and followed the guidelines for care and usage of laboratory animals of the Federal University of Rio Grande do Norte.

### 5.2 Virus injection of optogenetic constructs

Approximately 4 weeks prior to experiments using optogenetic stimulation, mice were injected with viral constructs of different opsins coupled to either channelrhodopsin2 (ChR2) or Archaerhodopsin3.0 (eArch3.0). ChR2 is a light activated cation permeable channel (Boyden et al., 2005) for membrane depolarization, while eArch3.0 is a green light activated outward proton (H+) pump (Chow et al., 2010) for membrane hyperpolarization. ChR2 constructs used were: rAAV5/CaMKII*α*-hChR2(H134R)- EYFP (Vector core, at a concentration of 4×10^12^ virus molecules – vm/ml) and a cre-dependent (double-floxed inverted open reading frame; DIO) ChR2 construct, AAV2/9.EF1a.DIO.hChR2(H134)- eYFP-WPRE-hGH (Vector core, at 1×10^13^ vm/ml). The eArch3.0 used was rAAV5/CamK2a-eArch3.0-eYFP (Vector core, at 2.5×10^12^ vm/ml). In detail, mice were anesthetized with ketamine-xylazine at 90/6 mg/kg intraperitoneal (i.p.). If required, additional ketamine was re-administered (as half the dose of the previous injection, i.e. 45 mg/kg and 22.5 mg/kg) during surgery. The mouse was mounted into a stereotaxic device while resting on a heating block at 37°C. Eye gel (dexpanthenol) was applied to avoid drying of eyes during surgery. The head was wiped with polyvidone-iodine (10%) to avoid infections. The skin was anesthetized with lidocaine hydrochloride 3% before a straight incision was made. After the incision, hydrogen peroxide 3% was applied onto the exposed skull to remove the connective tissue and to visualize sutures.

The DCN coordinates were taken from Franklin and Paxinos (Franklin et al., 2008). Specifically, we used −6.1mm anteroposterior (AP), −2.3mm mediolateral (ML), and −4.3mm and −3.8mm dorsoventral (DV, two steps). For each animal, those coordinates were corrected by multiplying by the normalized bregma-lambda distance (mouse’s bregma-lambda in mm divided by 4.2 – the average bregma-lambda distance from the mice used in Paxinos and Franklin’s atlas), to account for head size differences. Additionally, the vertical distance between the bregma and the point in the skull at the AP and ML coordinate was subtracted from the DV coordinate, so that the DCN can be reached using the brain surface as reference. A small mark was made at the AP and ML coordinates and a small hole was carefully drilled with a dental microdrill (Beavers Dental, Morrisburg, Canada). Next pre-aliquoted virus (20% for Cre-dependent, 30% for CaMKII*α*-dependent vectors) was rapidly thawed and withdrawn into a 10*μ*l Nanofil syringe with a 34-gauge removable needle, at the speed of 1.5 *μ*l/min using a infusion pump (Chemyx NanoJet). The syringe was lowered into the DCN to the deepest DV coordinate, and 0.75 *μ*l of virus was slowly infused at 0.15 *μ*l/min and the needle was kept in place for five minutes to allow for full diffuse of virus, then retracted to the second, more superficial DV coordinate for a second infusion (0.75 *μ*l) of virus and the needle kept in place for 10 minutes, before carefully removed. Some animals received bilateral injections. Next the skin was sutured, lidocaine hydrochloride 3% was applied over the suture and 200*μ*l of saline was injected subdermal in the back for rehydration. The animal was removed from the stereotaxic frame and placed under a red heat lamp and monitored until recovering from anesthesia. Some initial experiments used fluorescent retrobeads (Green fluorescent retrobeads, Lumafluore) to establish the appropriate coordinates of injection. The benefits of initially using retrobeads, compared to viral injections of optogenetic material during optimization of experimental procedures, are 1) the fluorescent liquid can be readily seen withdrawn into the microsyringe with the naked eye (compared to a minute volumes of transparent viral solution that sometimes fails to be withdrawn due to technical issues), and 2) animals can be sacrificed after only a few days (compared to waiting 2-4 weeks for viral expression) to confirm the appropriate location of fluorescent signal.

### 5.3 Sound calibration and sound stimulation

As different sound devices can have inherent shifts in unit level, and thereby in the signal generation, the sound card was initially calibrated using an oscilloscope. A 10kHz sine wave of 1V amplitude was written to the card, and the sound card output amplification factor was recorded as 1 divided by the amplitude of the output signal. All sound signals were multiplied by the output amplification factor before being written to the card. We connected the sound card output to the sound card input, and a 1V 10kHz sine wave was played and recorded. The input amplification factor was measured as 1 divided by the amplitude of the recorded signal, and signals read from the board were multiplied by it before any further processing. A loudspeaker (Super tweeter ST400 trio, Selenium Pro) was calibrated using a microphone (4939-A-011, Brüel and Kjær, Denmark) 4.5-10 cm in front of the speaker. Sound pulses (2s duration) were generated at the desired frequency bands with logarithmically decreasing amplification factors (voltage output to the speaker) and simultaneously recorded using a personal computer, and the power spectral density (PSD) of the recorded signal was calculated using a Hann window with no overlap. Root mean square (RMS) was calculated as

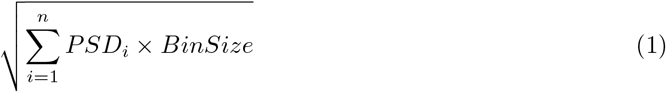

where *PSD* is a 1×*n* array and *BinSize* is the spectral resolution. The intensity in decibels sound pressure level (dBSPL) was calculated as

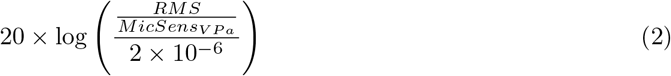

where MicSens_vpa_ is the microphone sensitivity in V/Pa, 0.004236V/Pa for our microphone. All data was saved to disk and loaded to provide the correct amplification factors for each sound intensity used for sound stimulation. The frequency band generated corresponds to the frequency band of greatest power in the signal spectrum, with border frequencies strongly attenuated (Supplementary Figure S1). Sound calibrations were routinely repeated before every beginning of an experimental group. The full sound calibration tests 300 amplification factors for each frequency band, providing 0.5 dBSPL precision. All hardware described here are outlined in Supplementary Table S1.

Sound stimulation consisted of sound pulses of gaussian white noise filtered from 5 to 15kHz, intensity of 80dBSPL and duration of 3ms, presented at 10Hz (3ms of sound pulse followed by 97ms of silence), repeated for 5 blocks of 200 pulses.

### 5.4 Light calibration and optogenetic stimulation

Light intensity calibration was performed before each experiment. Optic fibers of 200*μ*m diameter were cleaned with lens cleaning tissue and ethanol (99.5%). The light intensity was measured and the laser lens position adjusted until light power at the tip of the fiber was 5-7mW/mm^2^ measured by an optical power meter (Thorlabs PM20). Light stimuli triggers were generated in Python and written to the sound card (USBPre2; Thomann GmbH, Burgebrach, Germany), in which the output was splitted and connected to the laser input and the data acquisition board. Light stimulus was delivered using a 473nm laser (for ChR2) and a 532nm laser (for eArch3.0). Laser stimuli consisted of 200 light pulses at intensity of 5-7mW/mm^2^ with 10ms duration, presented at 10Hz (10ms on and 90ms off) 473nm blue light for ChR2 experiments; and a pulse of a total of 20s of 543nm green light repeated in 5 blocks with 10s interval, so that green light was continuously on during each sound stimulation block for eArch3.0 experiments. For concomitant sound and blue light stimulation the light pulses were presented 4ms before the sound pulses, so that these are embedded in the light pulse.

### 5.5 Digital timestamps marks for sound and light stimulation

Digital timestamps markers use a combination of a sound card (USBPre 2) and an Arduino board (Arduino Due, Arduino, Italy), taking advantage of GNU/Linux audio real-time capabilities. One output of the sound card was connected to the sound amplifier (PM8004, Marantz, New Jersey, USA), and another output to an acquisition board (Open-ephys, Open-ephys.org) analog input with a diode as rectifier, conducting only positive voltages. In detail, the two outputs of the sound card are used for sound stimulation (channel 1) and for generating timestamp marks (channel 2). To test the temporal accuracy of the digital input of the acquisition system (Open ephys), we recorded 5000 square pulses delivered by both analog and digital inputs to the acquisition board. When compared to analog traces, we found 15% of the digital timestamps to be delayed by >150*μ*s, which is a jitter of 5% of the 3ms pulse width (Supplementary Figure S2A). To avoid jitter, analog square waves to mark stimulation timestamps were used. To avoid producing capacitive-like traces when using square pulses in a sound card, we used square waves of twice the stimulus duration, containing both positive and negative portions (Supplementary Figure S2B). In practice, channels carrying square waves are connected to a diode before connecting to the analog input of the acquisition board or to the laser, thereby only conducting the positive values (Supplementary Figure S2B-C). The resultant square waves have the same duration as the stimulus, since only the positive half is conducted (Supplementary Figure S2B) thereby channel 2 was used both as timestamp marker and as a trigger for light stimulation.

For experiments using sound synchronized with light stimulation three outputs would be required (one carrying the sound signal to a speaker, one carrying the sound square waves /timestamps and the third carrying square waves for light trigger/timestamps). Here a square wave of twice the length for light stimulation was used, so when simultaneous sound and light stimulation is required, the sound pulse is written to channel 1 while the sum of the sound and light square waves are written to channel 2 (Supplementary Figure S2D). Thereby channel 2 triggers the laser (amplitude >3.3V), as well as provides edges for timestamp detection.

### 5.6 In vivo units recording

Animals were anesthetized with ketamine-xylazine (90/6 mg/kg i.p.) and an additional injection of (ketamine 45 mg/kg) if surgery required. The anesthetized mouse was placed on an electric thermal pad (37°C) and fixed into a stereotaxic frame with ear bars holding in front of and slightly above ears, on the temporal bone, to not block the ear canals. The skin over the vertex was removed and hydrogen peroxide (3%) was applied on the skull to visualize sutures. All coordinates were corrected as for the virus injection procedure. Next, three small holes were drilled: at AP=- 6.1mm ML=-2.3mm (left DCN, for probe placement); at AP=-6.1mm ML=2mm, (for optic fiber placement); and at AP=-2mm ML=1mm (for reference). Next, a micro screw was fixed in the reference coordinate using Polymethyl methacrylate. The optic fiber was inserted into the brain using a micromanipulator positioned in a 45° angle to a dept of 5.58mm, ending 0.5mm away from the DCN. This angle avoids perturbing auditory pathways and gives appropriate space for the insertion of the 16 channel single shank recording electrode. The recording electrode (single shank, 16 channels, 50*μ*m channel spacing, 177*μ*m^2^ recording site, 5mm length, Neuronexus) was dipped in fluorescent dye (1,1’-dioctadecyl-3,3,3’,3’-tetra -methylindocarbocyanine perchlorate; DiI, Invitrogen) for 10 minutes before the procedure to visualize electrode placement post-hoc. Channels are depict in figures as channel 1 being most dorsal and channel 16 most ventral. Three different recording depths were used (electrode tip at DV=4.0mm, 4.3mm or 4.5mm). Unit responses were recorded under sound and/or light stimulation, with modalities presented at randomized order. Data acquisition was done using a headstage (Intan RHA2116 or Intan RHD2132) connected to a data acquisition board (Intan RHA2000 or Open-ephys), at a sampling rate of 25kHz (for experiments with Intan RHA) or 30 kHz (for experiments with Open-Ephys). The headstage reference and ground were separated in the headstage, then ground was connected to the system ground and reference was connected to the reference screw. Responses were visualized using Open-ephys graphical user interface (Siegle et al., 2017) (GUI). At the end of recordings animals were sacrificed for histology.

### 5.7 Unit analysis

Spikes were detected and clustered with 4th order butterworth digital bandpass filter (300 to 7500 Hz), negative spikes detected using a threshold from 2-4.5× standard deviation (SD), waveforms of 2ms around the detected negative peak, and 3 features per channel. Peristimulus-time histograms (PSTHs) were calculated by counting occurrence of spikes in a time window of 100 ms around each TTL (50ms before and 50 ms after the TTL) and presented as mean number of spikes per time, where each bin corresponds to 1ms. Units were classified as responding units as described by Parras et al (Parras et al., 2017). In brief, 1000 PSTHs are generated with random values in a poisson distribution, with *lambda* equals to the mean of values from the negative portion of the unit PSTH (baseline). Then, for real and simulated PSTHs, the mean of the negative PSTH values (baseline) is subtracted from the mean of the positive PSTH values (response), resulting in a baseline-corrected spike count. Finally, the p-value is calculated as

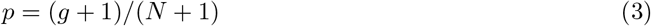

where *g* is the number of simulated histograms with corrected spike count bigger than the real unit spike count, and *N* is the total number of simulated histograms, which here is the number of trials presented at that unit recording. A cell was classified as responsive for a stimulation if the resulting p-value was <0.05. Cells were classified as responsive to *sound only, light only, sound*+*light only* or *sound and light*. Additionally, cells responding to sound stimulation were classified as *light-masked* when they respond to sound stimulation but do not respond to sound+light stimulation. Spike rate was calculated as spike events per second along all the recording (including the stimulation period). The threshold of 9 Hz was considered to separate between slow- and fast-spiking neurons, since ∼88% of neurons had firing rate < 6.42Hz and the remaining ∼12% had firing rate > 9.24Hz. Student’s t-test, two-tailed, unequal variance was applied to compare firing rate between neurons, and all firing rate values are represented as frequency ± standard error of the mean (s.e.m).

### 5.8 Auditory brainstem responses

Auditory brainstem responses (ABRs) were extracted from the same extracellular unit recordings. Data was filtered (4th order butterworth digital bandpass filter from 500 to 1500Hz), sliced (3ms before and 12ms after each sound pulse) and averaged. ABR peaks were detected as a positive value one standard deviation (SD) above the mean, larger than the previous value, and larger or equal the next value. ABR peak values and latencies were then grouped by sound or light stimulation (see Figure 8).

### 5.9 Histology

Intracardial perfusion was carried out by deeply anesthetizing mice with ketamine/xylazine (180/12 mg/kg). Animals were fixed in a polystyrene plate, and a horizontal incision was made in the skin at the level of the diaphragm. Thoracic cavity was open by cutting the ribs laterally and the sternal medially. A 30G needle was inserted into the left ventricle for perfusion with cold phosphate buffered saline (PBS) and an incision was made in the right atrium to allow for out-flow. In total, 20-30ml of cold PBS followed by 20-30ml of fixative (4% paraformaldehyde in 0.1M phosphate buffer; pH 7.4) was used. Next, the brain was dissected and stored in 4% paraformaldehyde overnight. For free-floating vibratome (OTS-4000, EMS, Hatfield) sections the brain was stored in PBS before slicing; and for cryostat sections, the brain was kept in PBS with 30% sucrose until dehydrated (visualized by the brain sinking to the bottom of the solution), and frozen using isopentane at −60°C. Horizontal sections (120*μ*m thick) of the brainstem, containing the DCN, were collected on glass slides and kept dark until examination of fluorescent expression by neurons. Cell nuclei were stained with 4’,6-diamidino-2-Phenylindole (DAPI) (Sigma) to visualize cell layers and borders of the DCN and VCN. Expression of optogenetic proteins was visualized by detection of genetically expressed eYFP. Images were collected using Zeiss Observer Z1 fluorescence microscope or a Zeiss Examiner Z1 confocal microscope. The objectives N-Achroplan 5x/0.15; N-Achroplan 10x/0.25; Plan-Apochromat 20x/0.8; and Plan-Neochromat 40x/0.75 were used. Images were collected using AxioVision and Zen software, respectively, and edited for brightness and contrast in ImageJ (NIH, Schneider et al., 2012).

### 5.10 CLARITY

The CLARITY procedure followed standard protocol and was previously describe for another brain region (Hilscher et al., 2017). Data from the auditory brainstem was collected during the same experiment as previously published (Hilscher et al., 2017) while the video attached here (Supplementary Video S1) was compiled specifically for the cochlear nucleus containing region.

### 5.11 Software availability

Our optogenetic and sound stimulation uses open systems and free open-source software. Recordings were done using Open-ephys GUI (Siegle et al., 2017). Calculations were done using Scipy (Jones et al., 2001) and Numpy (Van Der Walt et al., 2011), and all plots were produced using Matplotlib v2.2.4 (Caswell et al., 2019; Hunter, 2007). Spikes were detected and clustered using Klusta, and visual inspection was performed using Phy (Rossant et al., 2016). All scripts used for stimulation control and data analysis are available at https://gitlab.com/malfatti/SciScripts.

## Author Contributions

TM and BCB collected and analyzed electrophysiological and histological data under supervision of RNL and KEL; MMH and SJE analyzed CLARITY data under supervision of KK; TM, BCB and KEL wrote the manuscript with inputs from MMH and RNL.

## Acknowledgments

We would like to thank Dr. Helton Maia, Dr. George Nascimento and Dr. Joaõ Bacelo for technical advice.

## Data Availability Statement

The datasets generated and/or analyzed in the current study are available on request.

