## Supplemental Figures, Tables and Supplementary Video legend for "Using cortical neuron markers to target cells in the dorsal cochlear nucleus"

### 1 Supplementary figures

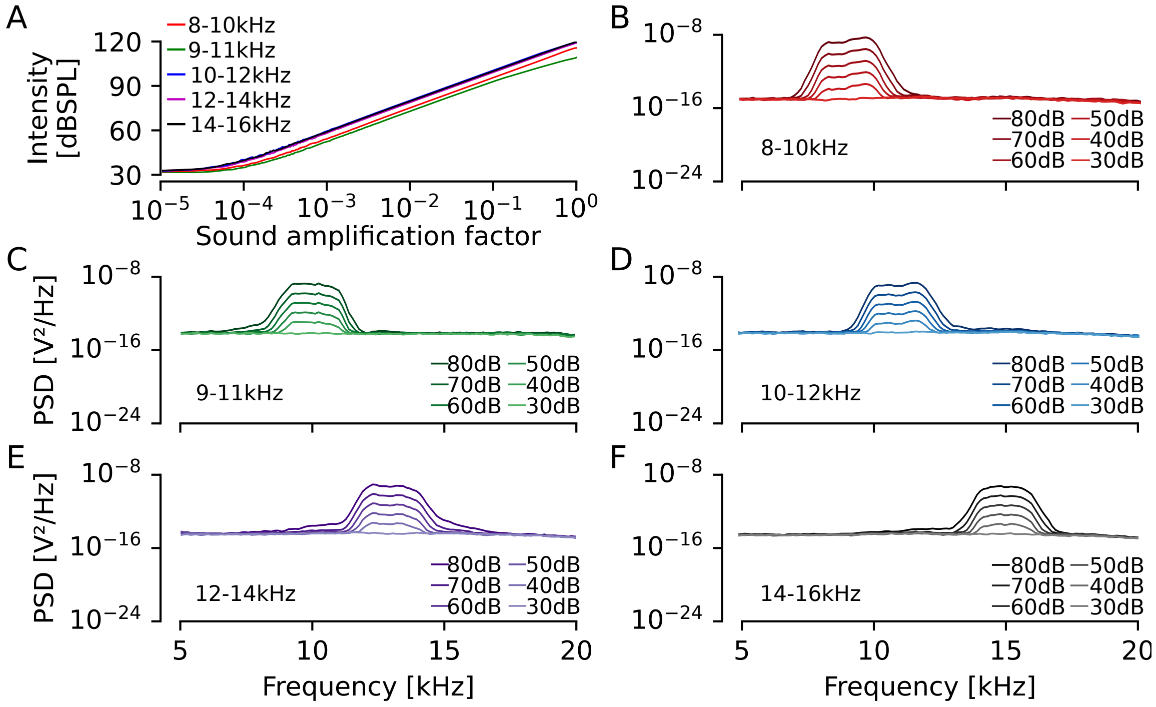


Figure 1: **Outline of sound calibration properties**. A) Intensity (dBSPL) in function of voltage (RMS) applied to the speaker. B-F) Power spectral density of six decreasing sound intensities for each frequency band tested.

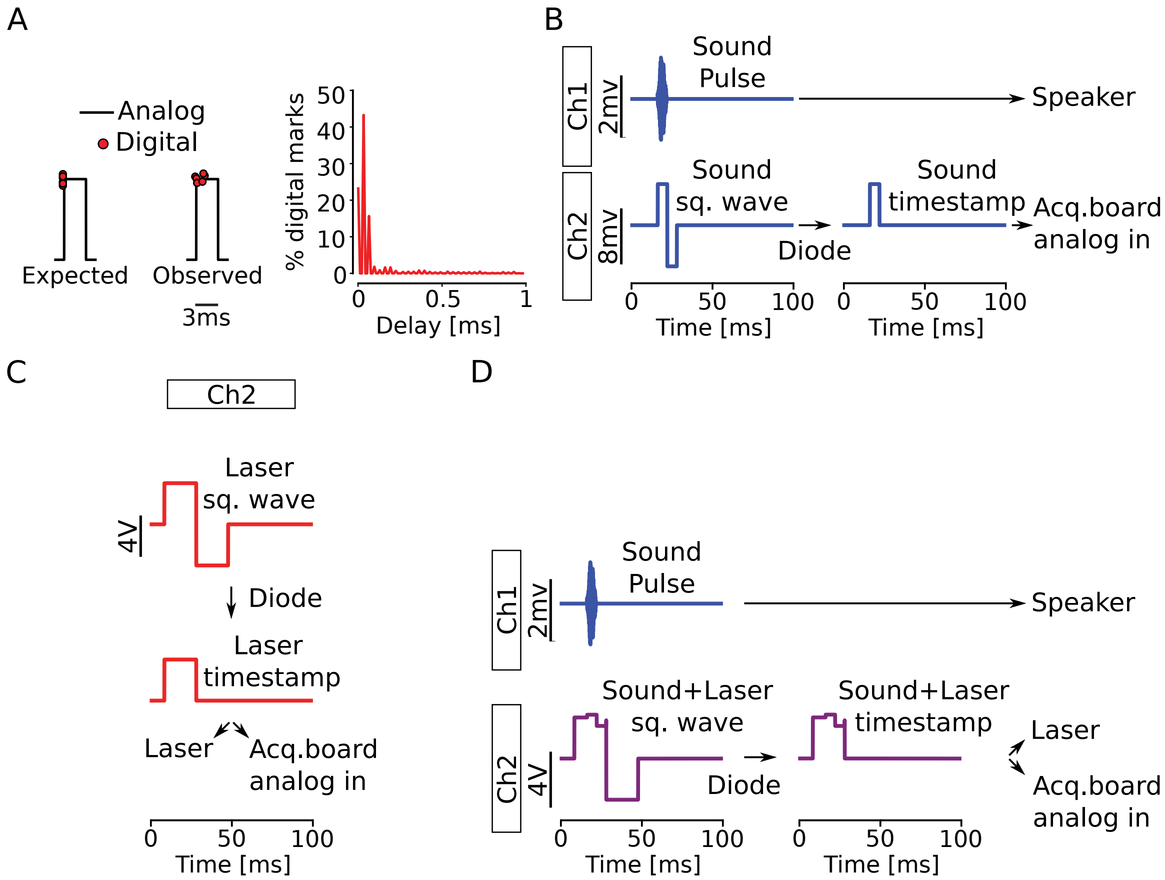


Figure 2: **Schematics of sound and light triggers combined with digital time stamps using a two channel sound card.** A) Schematic showing analog pulses and a small jitter of digital detected timestamps. *Right;* Histogram showing percentage of recorded pulses and delay to analog edge. B) Illustration of channel 1 producing a sound wave to the speaker; channel 2 producing the corresponding square wave with the positive portion with the same duration as the sound pulse (gray shading). Using a diode cuts the negative portion and can be detected as the sound timestamp. C) Channel 2 writes the light square wave and the positive portion is split to provide a laser trigger and the acquisition board analog input indicates the light timestamp. D) Channel 1 as 'B', and channel 2 illustrates the sum of light and sound square waves. As in 'C', both timestamps can be extracted from the signal recorded from the analog input of the acquisition board.

#### 2 Supplementary Tables

Table 1: List of hardware used for measurements, recording and stimulation; and software used to control the hardware. ADC = analog-digital converter; DAC = digital-analog converter.

| **Hardware** | **Company** | **Specifications** |
|---|---|---|
| Acquisition board | Open-ephys | v2.2, Opal Kelly XEM6010-LX150, 30KHz sampling rate, 16bit, 8 ADCs, 8 DACs |
| Computer | HP Z220 | Intel Xeon 8-core 3.6GHz, 16GB RAM |
| Headstage | Intan RHD2132 | 16 unipolar channels |
| Lasers | CNI | MBL-III-473 (blue) and MGL-III-532 (green) |
| Microcontroller | Arduino Due | 54 digital pins, 12 12bit ADCs, 2 12bit DACs |
| Microphone | Brüel and Kjær 4939-A-011 | 1/4 free-field microphone, sensitivity of 4.23643mV/Pa |
| Multichannel electrode | NeuroNexus | Single shank, 16 channels, 50µm channel spacing, 177µm$^2$ recording site, 5mm length |
| Sound card | Sound Devices USBPre2 | 192KHz sampling rate, 24bit ADC |
| Sound amplifier | Marantz PM8004 | |
| Speakers | Selenium Trio ST400 | |

Table 2: Number of recorded units, separated by group, responsiveness, firing rate and direction of modulation (down arrow decrease and up arrow increase firing rate upon concomitant sound and light stimulation comparing to sound alone).

| | n | **Resp.** | Resp. (Low FR) | Resp. (High FR) | **Non-resp.** | Non-resp. (Low FR) | Non-resp. (High FR) |
|---|---|---|---|---|---|---|---|
| CaMKIIa-ChR2 | 224 | 76 | 65 (↓ 34/↑ 31) | 11 (↓ 7/↑ 4) | 148 | 78 (↓ 26/↑ 52) | 70 (↓ 37/↑ 33) |
| CaMKIIa-eArch3.0 | 86 | 17 | 16 (↓ 4/↑ 12) | 1 (↓ 1/↑ 0) | 69 | 46 (↓ 31/↑ 15) | 23 (↓ 12/↑ 11) |
| Chrna2-cre/DIO-ChR2 | 76 | 15 | 14 (↓ 9/↑ 5) | 1 (↓ 0/↑ 1) | 61 | 30 (↓ 14/↑ 16) | 31 (↓ 13/↑ 18) |

#### 3 Supplementary Video

**Video S1.** Tomato+ cells (white) of Chrna2-Cre/R26tom mice visualized across sections corresponding to the auditory brainstem. A series of images from adult (2 months old) Chrna2-Cre/R26tom mouse brainstem (coronal slice, 1300 µm thickness) after CLARITY processing showing the ventral and dorsal portions of the cochlear nucleus.
